# *In vivo* validation of late-onset Alzheimer’s disease genetic risk factors

**DOI:** 10.1101/2023.12.21.572849

**Authors:** Michael Sasner, Christoph Preuss, Ravi S. Pandey, Asli Uyar, Dylan Garceau, Kevin P. Kotredes, Harriet Williams, Adrian L. Oblak, Peter Bor-Chian Lin, Bridget Perkins, Disha Soni, Cindy Ingraham, Audrey Lee-Gosselin, Bruce T. Lamb, Gareth R. Howell, Gregory W. Carter

## Abstract

**Introduction:** Genome-wide association studies have identified over 70 genetic loci associated with late-onset Alzheimer’s disease (LOAD), but few candidate polymorphisms have been functionally assessed for disease relevance and mechanism of action.

**Methods:** Candidate genetic risk variants were informatically prioritized and individually engineered into a LOAD-sensitized mouse model that carries the AD risk variants APOE4 and Trem2*R47H. Potential disease relevance of each model was assessed by comparing brain transcriptomes measured with the Nanostring Mouse AD Panel at 4 and 12 months of age with human study cohorts.

**Results:** We created new models for 11 coding and loss-of-function risk variants. Transcriptomic effects from multiple genetic variants recapitulated a variety of human gene expression patterns observed in LOAD study cohorts. Specific models matched to emerging molecular LOAD subtypes.

**Discussion:** These results provide an initial functionalization of 11 candidate risk variants and identify potential preclinical models for testing targeted therapeutics.

## Background

Alzheimer’s disease (AD) is the most common cause of dementia, with a growing clinical, financial, and social impact. An increasing body of evidence highlights the importance of genetic risk in AD (1-3). While a small percentage of AD cases are linked to causative, familial mutations in the amyloid precursor protein (APP) processing pathway, the vast majority of cases are late-onset AD (LOAD), have heterogeneous symptoms and etiology, and are associated with polygenic risk from a combination of low-risk, relatively common variants (4-6). Genome-wide association studies (GWAS) have identified numerous LOAD risk variants, but few have been experimentally validated, and physiological mechanisms have not been elucidated, even for the single strongest risk variant, the ε4 allele of *APOE* gene (4, 7). This is but one example (8) of the general problem of how to progress from the identification of genetic variants to functional impact of variants to getting to physiological disease mechanisms (9). Here we present a novel approach to assay the impact of individual polygenic risk factors using an *in vivo* approach.

While numerous potential therapeutics have shown promising results in transgenic mouse models of familial AD, few have advanced in clinical trials. This may result from numerous causes, but it is clear that one reason may be the lack of translational animal models available for preclinical studies (10-12). Almost all existing rodent models are based on causative mutations in proteins in the amyloid precursor protein (APP) processing pathway expressed in neurons. Most AD genetic risk resides in genes mainly expressed in microglia and other non-neuronal cell types, as recently reviewed (5, 13, 14), indicating that complex cellular interactions play a causative role in disease etiology. While *in vitro* systems have been shown to have value, more relevant *in vivo* models are necessary to understand these cell-cell interactions (15). In particular, animal models are required to study the early and progressive stages of pathology, which are not accessible in clinical studies but are critical to understand disease mechanisms so as to better target novel therapeutic approaches.

The MODEL-AD consortium was established to create and characterize translationally relevant mouse models of LOAD, and to set up protocols for preclinical testing in these new models (16). In this study we provide an overview of novel mouse models expressing human risk variants. Variants were introduced using a knock-in approach to avoid known issues with transgenic models (11, 17-19). To potentially enhance disease-relevant outcomes, variants were created on a more LOAD-susceptible genetic background expressing humanized *APOE* with the ε4 variant and the R47H mutation in *Trem2*, two of the strongest genetic risk factors for LOAD (20). The effects of each variant were assessed by gene expression changes in aging male and female brains using a newly developed transcriptomics panel (21), representing key LOAD-associated changes in clinical AD samples (22). This allowed us to functionalize GWAS variants with small but significant increases in disease risk and avoided a reliance on amyloid deposition or cognitive assays, which have not proven to translate to clinical studies.

## Methods

### Late-onset AD risk variant prioritization

Prioritization and construction of the APOE and TREM2 variants in the LOAD1 strain were previously discussed (20). Late-onset variants were selected based on human genetic association, predicted pathogenicity, conservation with mouse homolog, and allele frequency. We further prioritized based on diversity in predicted function to maximize our exploration of potential LOAD biology. Determining specific variants was primarily limited by the rarity of strong coding candidates (e.g. nonsynonymous, stop-gain) and strict mouse sequence homology that required the same SNP be engineered into mice. This led to a mix of variants at high-confidence GWAS loci, functional candidates, and exploratory variants. Exome sequencing from the Alzheimer’s Disease Sequencing Project (ADSP) was initially used to identify specific variants at loci (23), buttressed by summary data at NIAGADS (https://www.niagads.org/genomics/app). All variants are annotated as “ADSP Variants” that passed NIAGADS quality control checks (https://www.niagads.org/genomics/app).

ABCA7*A1527G (rs3752246) is the most common of multiple predicted loss-of-function variants associated with increased LOAD risk at the *ABCA7* locus (24, 25). The SORL1*A528T (rs2298813) variant is among candidates in the *SORL1* gene and likely involved in retromer function (26); deficits in retromer-dependent endosomal recycling have been implicated as causal in AD (27-29). The SNX1*D465N (rs1802376) variant locus is associated with AD (24) and *SNX1* is involved in retromer function relevant to LOAD (30). PLCG2*M28L (rs61749044) has been associated with LOAD [https://www.biorxiv.org/content/10.1101/2020.05.19.104216v1, (24, 31) and Plcg2 is a key protein in microglial activation in response to AD pathology (32). The SHC2*V433M (rs61749990) variant was identified in ADSP exomes and has been associated with neurodegeneration and neuron loss (33, 34). SLC6A17*P61P (rs41281364) reduces gene expression in the brain (gtexportal.org/home/gene/SLC6A17), and its reduction is also associated with LOAD (agora.adknowledgeportal.org/genes/ENSG00000197106). Rare variants have been associated with neurological phenotypes (35, 36). The CLASP2*L163P (rs61738888) variant has been associated with neurodegeneration from meta-analysis (37). The MTMR4*V297G (rs2302189) variant has been linked to cognitive function (38, 39). Predicted *CEACAM1* loss-of-function variants had a high disease burden in ADSP exome sequencing data (SKAT-O Bonferroni-adjusted p = 7.47 x 10-7) and the gene was associated with AD-related traits in a model of mouse genetic variability (40). The common MTHFR*677C>T (rs1801133) has been associated with increased risk for LOAD and other age-related disorders (41, 42). To explore a copy-number variant linked to vascular function, we used an existing MEOX2 knockout based on an association with Alzheimer’s disease (43) that may be related to the gene’s role in neurovascular health (44). This variant was assessed in a heterozygous state due to non-viability of the homozygote.

### Model development

All experiments were approved by the Animal Care and Use Committee at The Jackson Laboratory. Mice were bred in the mouse facility at The Jackson Laboratory and maintained in a 12/12-h light/dark cycle, consisting of 12 h-ON 7 am-7 pm, followed by 12 h-OFF. Room temperatures are maintained at 18–24LC (65–75LF) with 40–60% humidity. All mice were housed in positive, individually ventilated cages (PIV). Standard autoclaved 6% fat diet (Purina Lab Diet 5K52) was available to the mice ad-lib, as was water with acidity regulated from pH 2.5–3.0.

Novel mouse alleles were generated using direct delivery of CRISPR-Cas9 reagents to LOAD1 (JAX #28709)(20) mouse zygotes. Analysis of genomic DNA sequence surrounding the target region, using the Benchling (www.benchling.com) guide RNA design tool, identified appropriate gRNA sequences with a suitable target endonuclease site.

*Streptococcus pyogenes* Cas9 (SpCas9) V3 protein and gRNA were purchased as part of the Alt-R CRISPR-Cas9 system using the crRNA:tracrRNA duplex format as the gRNA species (IDT, USA). Alt-R CRISPR-Cas9 crRNAs (Product# 1072532, IDT, USA) were synthesized using the gRNA sequences specified in the DESIGN section and hybridized with the Alt-R tracrRNA (Product# 1072534, IDT, USA) as per manufacturer’s instructions. Plasmid or oligonucleotide constructs were synthesized by Genscript.

See supplemental Table 1 for CRISPR reagents.

To prepare the gene editing reagent for electroporation, SpCas9:gRNA Ribonucleoprotein (RNP) complexes were formed by incubating AltR-SpCas9 V3 (Product#1081059, IDT, USA) and gRNA duplexes for 20 minutes at room temperature in embryo tested TE buffer (pH 7.5). The SpCas9 protein and gRNA duplex were at 833 ng/ul and 389 ng/ul, respectively, during complex formation. Post RNP formation, the purified plasmid was added and the mixture spun at 14K RPM in a microcentrifuge. The supernatant was transferred to a clean tube and stored on ice until use in the embryo electroporation procedure. The final concentration of the gRNA, SpCas9 and plasmid components in the electroporation mixture were 600ng/ul, 500 ng/ul and 20 ng/ul, respectively.

Founders were selected that: were positive by short-range PCR assays; had appropriate sequence across the homology arm junctions; were negative for the plasmid backbone; and had correct sequence of the inserted construct.

Allele-specific genotyping protocols for all models are available on JAX Mice data sheets for each model.

Other models were obtained from the JAX mouse repository, see Table 1.

**TABLE 1:** Listing of gene loci, human risk variants and corresponding mouse alleles, allele type, and JAX ID of mouse models created. All models also contain a humanized APOE4 allele and a Trem2*R47H allele on the C57BL6/J background (“LOAD1”), which was used as a control.

| Locus | Allele (Human) | Allele (Mouse) | SNP | Allele Type | JAX # |
| --- | --- | --- | --- | --- | --- |
| Abca7 | A1527G | A1541G | rs3752246 | missense | 30283 |
| Ceacam1 | LOF variants | KO | --- | KO | 30673 |
| Clasp2 | L163P | L163P | rs61738888 | missense | 31944 |
| Meox2 | LOF variants | HET KO | --- | HET KO | 33770 |
| Mthfr | A222V (677C>T) | A262V | rs1801133 | missense | 30922 |
| Mtmr4 | V297G | V297G | rs2302189 | missense | 31950 |
| Plcg2 | M28L | M28L | rs61749044 | missense | 30674 |
| Shc2 | V577M | V433M | rs2298813 | missense | 31952 |
| Slc6a17 | P61P | P61P | rs41281364 | silent mutation | 31948 |
| Snx1 | D466N | D465N | rs1802376 | missense | 31942 |
| Sorl1 | A528T | A528T | rs41281364 | missense | 31940 |
| **Other models used:** |  |  |  |  |  |
| C57BL/6J |  |  |  |  | 664 |
| 5xFAD |  |  |  |  | 8730 |
| LOAD1 |  |  |  |  | 28709 |

### Brain Harvest at 4 months of age

Anesthetized and subsequently perfused animals were decapitated, and heads submerged quickly in cold 1X PBS. The brain was carefully removed from the skull, weighed, and divided midsagitally, into left and right hemispheres, using a brain matrix. The right hemisphere was quickly homogenized on ice and equally aliquoted into cryotubes for and transcriptomic analysis. Cryotubes were immediately snap-frozen on dry ice and stored long-term at -80LC.

### RNA Sample Extraction

Total RNA was extracted from snap-frozen right brain hemispheres using Trizol (Invitrogen, Carlsbad, CA). mRNA was purified from total RNA using biotin-tagged poly dT oligonucleotides and streptavidin-coated magnetic beads and quality was assessed using an Agilent Technologies 2100 Bioanalyzer (Agilent, Santa Clara, CA).

RNA-Sequencing Assay Library Preparation Sequencing libraries were constructed using TruSeq DNA V2 (Illumina, San Diego, CA) sample prep kits and quantified using qPCR (Kapa Biosystems, Wilmington, MA). The mRNA was fragmented, and double-stranded cDNA was generated by random priming. The ends of the fragmented DNA were converted into phosphorylated blunt ends. An ‘‘A’’ base was added to the 3’ ends. Illumina-specific adaptors were ligated to the DNA fragments. Using magnetic bead technology, the ligated fragments were size-selected and then a final PCR was performed to enrich the adapter-modified DNA fragments since only the DNA fragments with adaptors at both ends will amplify.

### RNA-Sequencing

Libraries were pooled and sequenced by the Genome Technologies core facility at The Jackson Laboratory. All samples were sequenced on Illumina HiSeq 4000 using HiSeq 3000/4000 SBS Kit reagents (Illumina), targeting 30 million read pairs per sample. Samples were split across multiple lanes when being run on the Illumina HiSeq, once the data was received the samples were concatenated to have a single file for paired-end analysis.

### RNA-Sequencing Data Processing

Sequence quality of reads was assessed using FastQC (v0.11.3, Babraham). Low-quality bases were trimmed from sequencing reads using Trimmomatic (v0.33; Bolger et al., 2014). After trimming, reads of length longer than 36 bases were retained. The average quality score was greater than 30 at each base position and sequencing depth was in range of 60–80 million reads. RNA-Seq sequencing reads from all samples were mapped to the mouse genome (version GRCm38.p6) using ultrafast RNA-Seq aligner STAR (v2.5.3; Dobin et al., 2013). To measure human APOE gene expression, we created a chimeric mouse genome by concatenating the human APOE gene sequence (human chromosome 19:44905754-44909393) into the mouse genome (GRCm38.p6) as a separate chromosome (referred to as chromosome 21 in chimeric mouse genome). Subsequently, we added gene annotation of the human APOE gene into the mouse gene annotation file. Additionally, we have also introduced annotation for novel Trem2 isoform in mouse gene annotation file (GTF file), that is identical to primary transcript but truncated exon2 by 119 bp from its start position(20). Afterward, a STAR index was built for this chimeric mouse genome sequence for alignment, then STAR aligner output coordinate-sorted BAM files for each sample mapped to the chimeric mouse genome using this index. Gene expression was quantified in two ways, to enable multiple analytical methods: transcripts per million (TPM) using RSEM (v1.2.31; Li and Dewey, 2011), and raw read counts using HTSeq-count (v0.8.0; Anders et al., 2015).

### NanoString transcriptomic analysis

The NanoString Mouse AD gene expression panel (21) was used for gene expression profiling on the nCounter platform (NanoString, Seattle, WA). Mouse NanoString gene expression data were collected from brain hemisphere homogenates at 4, 8 and 12 months of age for both sexes, from approximately six animals per group. The nSolver software was used for generating NanoString gene expression counts. Normalization was done by dividing counts within a lane by geometric mean of the designated housekeeping genes from the same lane. Next, normalized count values were log-transformed and corrected for potential batch effects using ComBat (45).

Next, we determined the effects of each factor (sex and genetic variants) by fitting a multiple regression model using the lm function in R as (46):

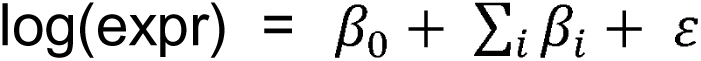

The sum is over sex (male), and all genetic variants (*5xFAD, LOAD1, Abca7*A1527G, Ceacam1KO, Mthfr*677C>T, Shc2*V433M, Slc6a17*P61P, Clasp2*L163P, Sorl1*A528T, Meox2 KO (HET), Snx1*D465N, Plcg2*M28L, Mtmr4*V297G*) used in this study. The log(expr) represents log-transformed normalized count from the NanoString gene expression panel (21). In this formulation, B6J was used as the control for the 5xFAD and LOAD1 mouse models, whereas LOAD1 served as controls for GWAS-based models in order to estimate the effects of individual variants. Separate models were run for each age cohort.

### Human AMP-AD Gene Co-expression Modules

Data for 30 human brain co-expression modules from the Accelerating Medicines Partnership for Alzheimer’s Disease (AMP-AD) studies were obtained from the Synapse data repository (https://www.synapse.org/#!Synapse:syn11932957/tables/; SynapseID: syn11932957). Briefly, Wan et al. (2020) (22) identified 30 human brain co-expression modules based on meta-analysis of differential gene expression from seven distinct brain regions in postmortem samples obtained from three independent LOAD cohorts (47-49). These 30 human AMP-AD modules were further classified into five distinct Consensus Clusters that describe the major functional alterations observed in human LOAD (21, 22).

### Human AD Subtypes

Milind et al. (50), integrated post-mortem brain co-expression data from the frontal cortex, temporal cortex, and hippocampus brain regions and stratified patients into different molecular subtypes based on molecular profiles in three independent human LOAD cohorts (ROS/MAP, Mount Sinai Brain Bank, and Mayo Clinic) (47-49). Two distinct LOAD subtypes were identified in the ROSMAP cohort, three LOAD subtypes were identified in the Mayo cohort, and two distinct LOAD subtypes were identified in the MSBB cohort. Similar subtype results were observed in each cohort, with LOAD subtypes found to primarily differ in their inflammatory response based on differential expression analysis (50). Data for LOAD subtypes were obtained through AD Knowledge Portal (51) (https://www.synapse.org/#!Synapse:syn23660885).

### Mouse-human expression comparison

To assess the human disease relevance of LOAD risk variants in mice, we determined the extent to which changes due to genetic perturbations in mice matched those observed in human AD cases versus controls. For each mouse perturbation, we tested each of the 30 AMP-AD modules using mouse-human gene homologs and limited to the genes both present in the module and the NanoString Mouse AD Panel, which was designed to optimize coverage of these modules (21). Pearson’s correlations were computed for changes in gene expression (log fold change) across all module genes for human AD cases versus controls (22) against the effect of each mouse perturbation (β) as measured above (21, 46). We used the cor.test function in R as:

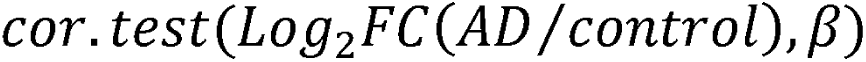

from which we obtained the correlation coefficient and the significance level (p-value) of the correlation for each perturbation-module pair. Log_2_FC values for human transcripts were obtained through the AD Knowledge Portal (51) (https://www.synapse.org/#!Synapse:syn14237651).

To determine the similarity of each mouse perturbation and the LOAD subtypes, we computed the Pearson’s correlation between gene expression changes (log fold change) in human AD subtype cases versus controls (50) and the effect of each mouse perturbation (β) across genes on the NanoString panel (21) using cor.test function in R as:

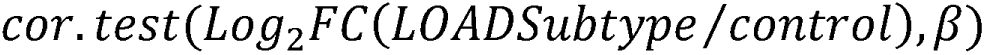

from which we obtained both the correlation coefficient and the significance level (p-value) of the correlation. Here, Log_2_FC(LOAD Subtype/control) represented the log-fold change in gene expression in each subtype versus control and the correlation spanned all homologous genes on the NanoString AD Mouse Panel.

We plotted the correlation results using the ggplot2 package in R. Framed circles were used to denote significant (p < 0.05) positive (blue) and negative (red) Pearson’s correlation coefficients. The color intensity and size of the circles were sized proportional to Pearson’s correlation coefficient.

### Functional enrichment analysis

Gene set enrichment analysis (GSEA) was used based on the method proposed by Subramanian, et. al (52) as implemented in the R Bioconductor package clusterProfiler (53) for the Reactome pathway library and Gene Ontology terms. Nanostring Mouse AD Panel genes (21) were ranked based on regression coefficients calculated for each factor and GSEA was performed on this ranked dataset. The use of GSEA ensured that pathway effects were assessed relative to the genes on the panel, as the panel was enriched for AD-relevant genes. Enrichment scores for all Reactome pathways and GO terms were computed to compare relative expression on the pathway level between each factor estimate from the regression models. We also performed Gene Ontology term enrichment analyses using enrichGO function in the clusterProfiler (53). Significance of pathways and GO terms were determined using false discovery rate (FDR) multiple testing correction (FDR adjusted p < 0.05).

## Results

### Validation of novel models

Sequence analysis demonstrated that the appropriate sequence variants had been established, see Supplemental Figure 1A. Quantification of transcript counts in homozygous LOAD models relative to littermate wild-type controls showed no significant differences in expression levels (Supplemental Figure 1B).

### LOAD associated risk variants showed age-dependent concordance with distinct human co-expression modules

We assess the relevance of each LOAD risk variant to the molecular changes observed in human disease (47-49, 54) by correlating the effect of each mouse perturbation (sex and genetic variants) with 30 human AMP-AD brain gene co-expression modules (22) using the NanoString Mouse AD Panel (21). We analyzed mouse NanoString data from brain hemispheres at different ages (4 and 12 months) to assess the correlation with human post-mortem co-expression modules as animals aged.

The amyloidogenic 5XFAD transgenic model exhibited significant positive correlations (p < 0.05) with several human co-expression modules in Consensus Cluster B enriched for immune-system related pathways at both four and 12 months but showed significant positive correlations (p < 0.05) with neurodegeneration associated human co-expression modules in Consensus Cluster C only at 12 months (Figure 2A-B). However, we did not observe significant positive correlations between effect of 5xFAD and human co-expression modules in Consensus Cluster A, D, and E, validating that the 5xFAD strain is primarily a model of amyloidosis and does not fully recapitulate late-onset AD changes.

**FIGURE 1:**
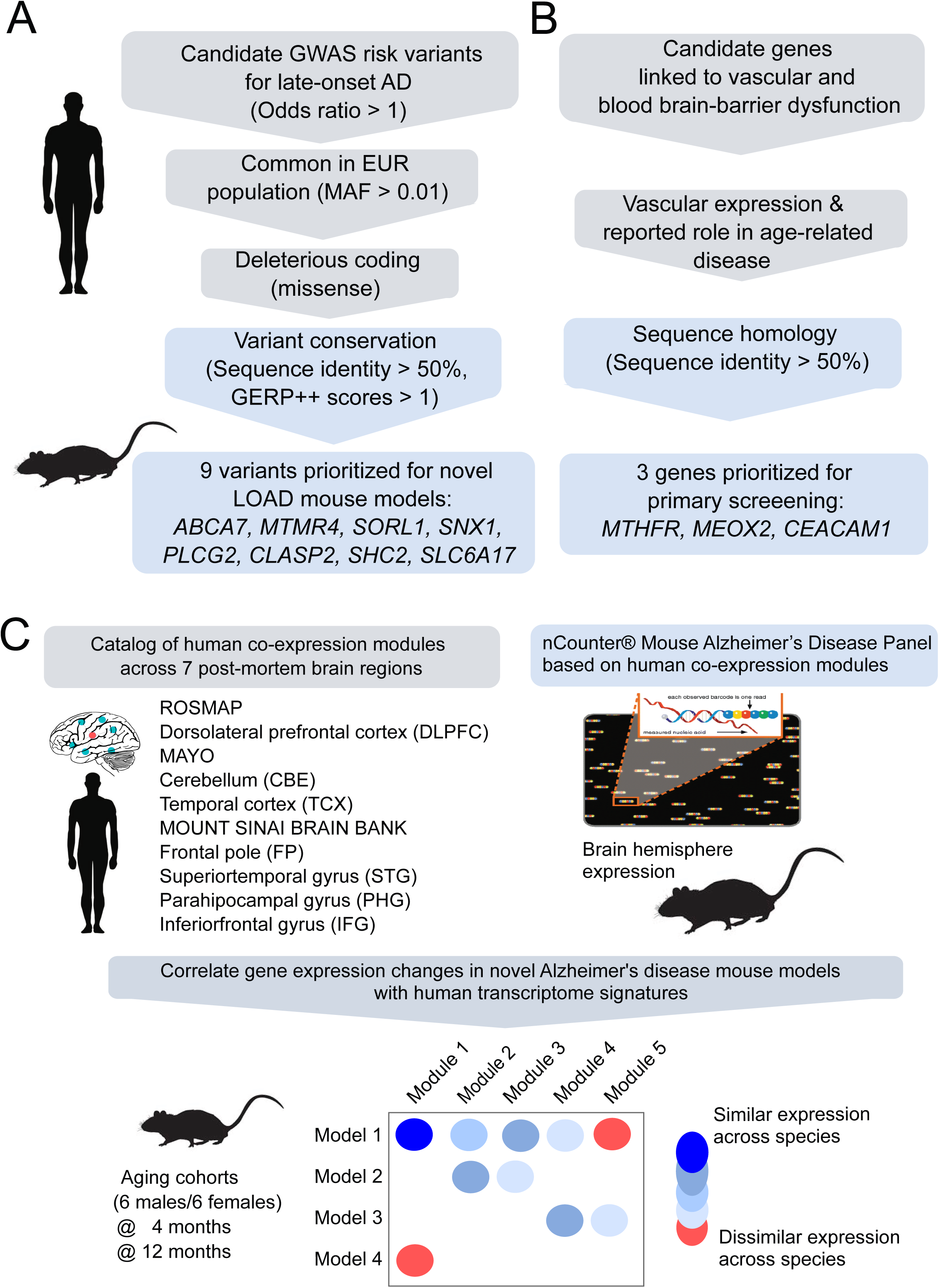
Strategy to prioritize loci and LOAD risk variants. Summary of strategies for variant selection for **(A)** late-onset Alzheimer’s disease and **(B)** neurovascular risk factors. **(C)** Gene expression analysis comparing human and mouse gene expression data to identify human LOAD modules that are altered by genetically engineered variants in mice.

**FIGURE 2:**
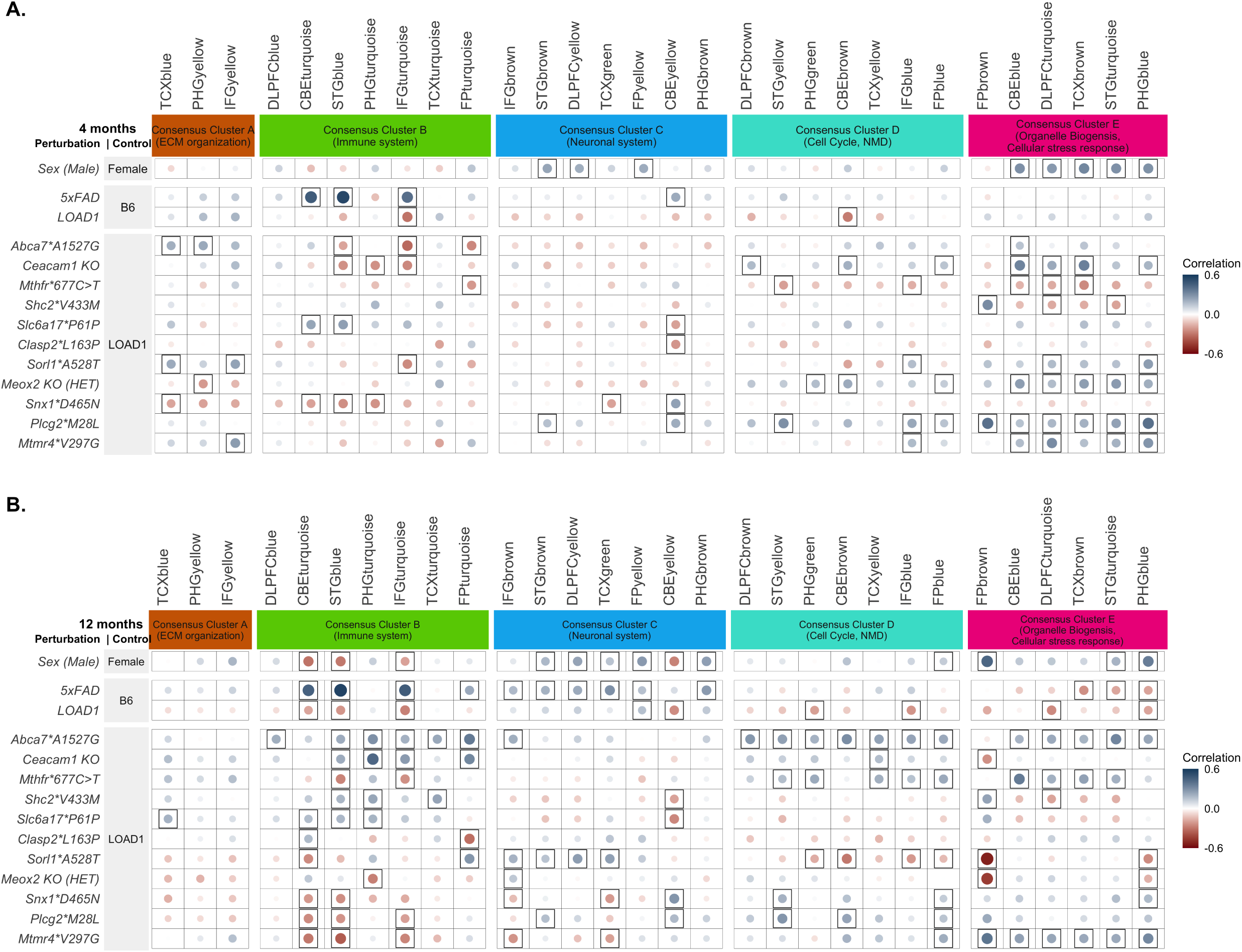
Correlation between LOAD associated risk variants and 30 human AMP-AD brain co-expression modules using the NanoString Mouse AD panel (A) Correlation between the effect of each mouse perturbation relative to the LOAD1 background in four-month-old mice and 30 human co-expression modules (22), also including the early-onset transgenic model 5XFAD and the LOAD1 background relative to C57BL/6J. The 30 human co-expression modules were grouped into five consensus clusters with similar gene content across the multiple studies and brain regions (22). Framed circles correspond to significant (p < 0.05) positive (blue) and negative (red) Pearson’s correlation coefficients, with size and color intensity proportional to the correlation. The effects of multiple LOAD risk variants in mice were positively correlated (p < 0.05) with cell cycle and myelination-associated modules in Consensus Cluster D and cellular stress-response associated modules in Consensus Cluster E. **(B)** Correlation between the effect of each mouse perturbation at 12 months and the 30 human co-expression modules. LOAD risk variants showed significant correlation with functionally distinct AMP-AD co-expression modules. The effects of *Abca7*A1527G*, *Shc2*V433M*, *Ceacam1 KO,* and *Slc6a17*P61P* in aged mice correlated with the immune modules in Consensus Cluster B, while the effects of *Sorl1*A528T and Plcg2*M28L* correlated with the neuronal modules in Consensus Cluster C.

At 4 months, among all LOAD risk variants, only *Slc6a17*P61P* showed significant positive correlations (p<0.05) with the immune related modules (Figure 2A). The *Abca7*A1527G, Sorl1*A528T*, and *Mtmr4*V297G* risk variants exhibited significant positive correlations (p<0.05) with extracellular matrix organization-related modules in Consensus Cluster A (Figure 2A). The *Ceacam1 KO*, *Plcg2*M28L, Meox2 KO(HET)*, and *Mtmr4*V297G* strains exhibited significant positive correlations (p<0.05) with cell cycle and myelination-associated modules in Consensus Cluster D and cellular stress-response associated modules in Consensus Cluster E (Figure 2A). *Abca7*A1527G and Sorl1*A528T* variants generated significant positive correlations (p<0.05) with cellular stress-response associated modules in Consensus Cluster E.

We observed more significant correlations between LOAD risk variants and human AMP-AD modules at 12 months for most strains. The *Abca7*A1527G* variant had the most pronounced correlations with LOAD expression changes, exhibiting significant positive correlations (p<0.05) with immune related modules in Consensus Cluster B, cell cycle and myelination-associated modules in Consensus Cluster D, and cellular stress-response associated modules in Consensus Cluster E (Figure 2B). The *Mthfr*677C>T* variant exhibited significant positive correlations (p<0.05) with cell cycle and myelination-associated modules in Consensus Cluster D and cellular stress-response associated modules in Consensus Cluster E (Figure 2B). *Sorl1*A528T* led to significant positive correlations (p<0.05) with several human co-expression modules in Consensus Cluster C enriched for neuronal related pathways (Figure 2B). The *Plcg2*M28L* variant had significant positive correlations (p<0.05) with human co-expression modules in Consensus Cluster C enriched for neuronal related pathways and with cell cycle and myelination-associated modules in Consensus Cluster D (Figure 2B). *Ceacam1 KO*, *Slc6a17*P61P, and Shc2*V433M* exhibited significant positive correlations (p<0.05) with human co-expression modules in Consensus Cluster B enriched for transcripts associated with immune related pathways in multiple brain regions, while *Clasp2*L163P* and *Sorl1*A528T* led to significant positive correlations (p<0.05) with human co-expression module in Consensus Cluster B enriched for immune related pathways in cerebellum and frontal pole brain region, respectively (Figure 2B). The *Mtmr4*V297G* variants exhibited significant positive correlations (p<0.05) with cell cycle and myelination-associated modules in Consensus Cluster D and cellular stress-response associated modules in Consensus Cluster E (Figure 2B). *Snx1*D465N* also exhibited significant positive correlation with cell-cycle and myelination-associated modules in Consensus Cluster D (Figure 2B).

Overall, we observed LOAD risk variants in mice showed concordance with distinct human co-expression modules, reflecting a different transcriptional response driven by each LOAD risk variant. The associations between LOAD risk variants and human gene co-expression modules increased with age. We note that models harboring late-onset AD risk variants exhibited significant positive correlation with human modules in Consensus Cluster A, D and E, which were not captured by 5XFAD strain, highlighting the importance of using LOAD risk variants to fully capture LOAD molecular pathologies.

We next assessed the similarities between variant effects in mice by comparing each model to all other models To identify LOAD risk variants driving similar transcriptional responses in mice, we performed correlation between regression coefficients calculated for each genetic variant at four and 12months. At four months, effects of the LOAD1 construct (*APOE4* and *TREM2*R47H*) were weakly and positively correlated with effect of 5xFAD transgene (p < 0.05), but this correlation diminished at 12 months (Figure 3A-B). Effects of LOAD1 were also significantly positively correlated (p<0.05) with *Sorl1*A528T* and *Mtmr4*V297G* at four months, but this correlation diminished by 12 months (Figure 3A-B). Effects of *Abca7*A157G* and *Ceacam1 KO* variants were weakly correlated at four months (p < 0.05), and this correlation increased at 12 months (Figure 3A-B). Effects of *Shc2*V433M* and *Slc6a17*P161P* variants were also significantly positively correlated at four months (p < 0.05) and become stronger with age (Figure 3A-B). Furthermore, effects of *Snx1*D465N*, *Plcg2*M28L,* and *Mtmr4*V297G* risk variants were significantly positively correlation (p < 0.05) at 12 months. Similarly, effects of the *Sorl1*A528T* and *Meox2 KO(HET)* variants were significantly positively correlated (p < 0.05) at 12 months (Figure 3A-B). In summary, we observed that LOAD risk variants generally increased in similarity with age, supporting an age-dependent role for these genetic factors. However, all strains did not converge on similar transcriptional responses, suggesting distinct mechanisms of influence on LOAD risk.

**FIGURE 3:**
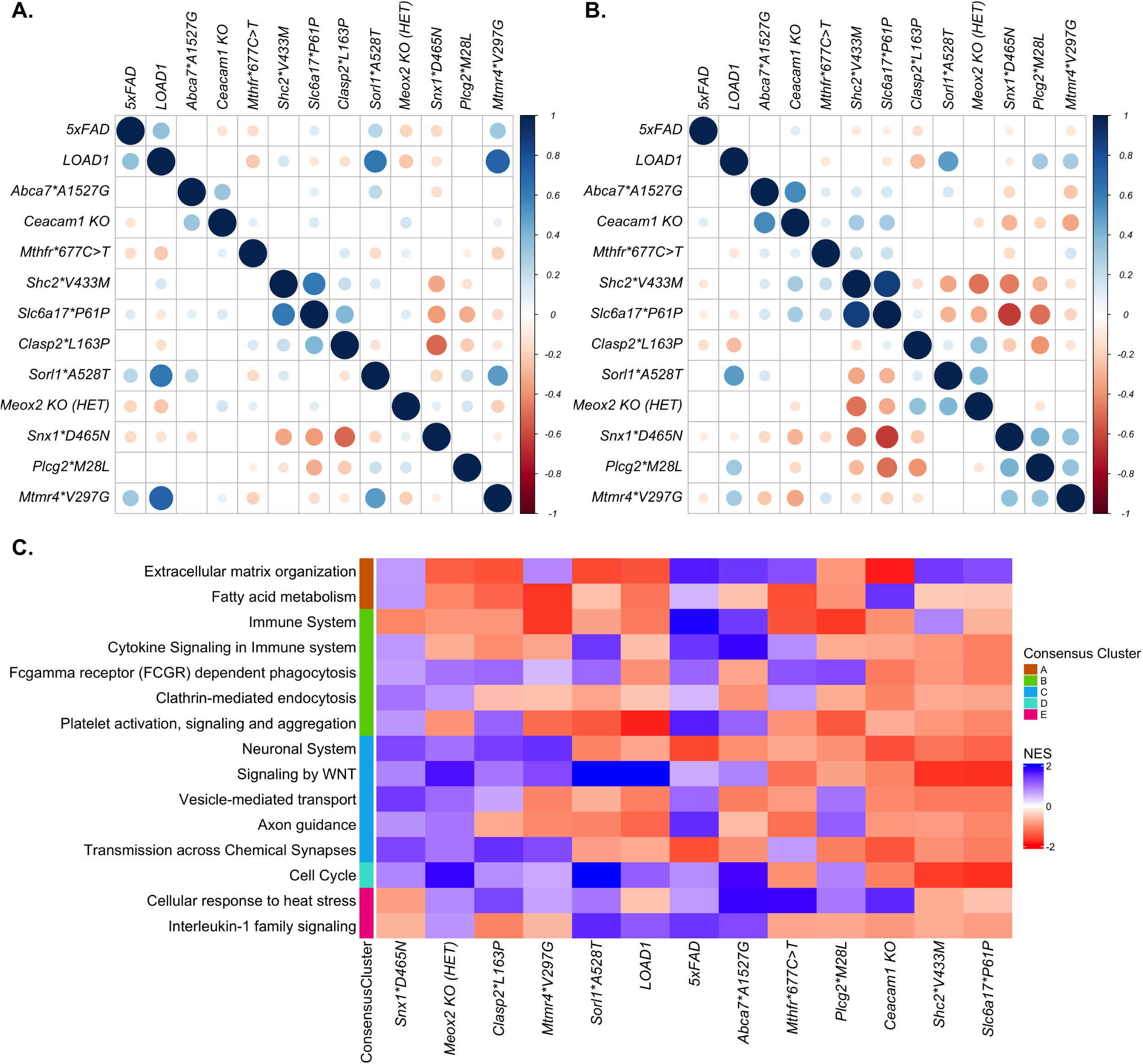
Correlation between effect of genetic variants and gene set enrichment analysis (A) Correlation between regression coefficients calculated for each genetic variant at four months. Color intensity and size of the circles are proportional to the Pearson correlation coefficient, with insignificant correlations (p > 0.05) left blank. **(B)** Correlation between regression coefficients calculated for each genetic variant at 12 months. The effects of *Snx1*D465N*, *Plcg2*M28L,* and *Mtmr4*V297G* risk variants in mice showed significantly positively correlation (p < 0.05) at 12 months **(C)**. Gene set enrichment analyses results of selected AD-associated pathways from Reactome library in the presence of each LOAD risk variants in mice. Enriched pathways are grouped by their overlap with functional annotations of human AMP-AD Consensus Clusters. Immune-related pathways had increased expression in the presence of multiple risk variants such as *Abca7*A1527G, Mthfr*677C>T, and Snx1*D465N,* while neuronal-associated pathways had reduced expression in the presence of risk variants such as *Abca7*A1527G, Mthfr*677C>T, Sorl1*A528T, Plcg2*M28L, Ceacam1 KO, Shc2*V433M, and Slc6a17*P161P*.

### Pathway alterations varied by LOAD genetic perturbation

To further elucidate the functional role of these LOAD risk variants in aged mice, we performed Gene Set Enrichment Analysis (GSEA) (52) for the Reactome pathway library for all 12 month samples. GSEA revealed upregulation of immune-related pathways in the presence of multiple risk variants such as *Abca7*A1527G, Mthfr*677C>T, and Snx1*D465N* (Figure 3C, Supplementary Table S2), while neuronal-associated pathways were downregulated in the presence of risk variants such as *Abca7*A1527G, Mthfr*677C>T, Sorl1*A528T, Plcg2*M28L, Ceacam1 KO, Shc2*V433M, and Slc6a17*P161P* (Figure 3C, Supplementary Table S2). Extracellular matrix organization pathway was downregulated in risk variants such as *Sorl1*A528T, Clasp2*L163P, Meox2 KO(HET)* and *LOAD1* but upregulated in the presence of risk variant such as *Abca7*A1527G, Snx1*D465N and Mthfr*677C>T* (Figure 3C, Supplementary Table S2). Cell cycle pathway was downregulated in the presence of *Mthfr*677C>T, Ceacam1 KO, Shc2*V433M, and Slc6a17*P161P,* while upregulated in the presence of other risk variants such as *Abca7*A1527G, Clasp2*L163P, Meox2 KO(HET),* and *Sorl1*A528T* (Figure 3C, Supplementary Table S2). Cellular response to heat stress pathway were downregulated in the presence of *Snx1*D465N*, *Shc2*V433M, and Slc6a17*P161P,* but upregulated in the presence of risk variants such as *Abca7*A1527G, Mthfr*677C>T, Sorl1*A528T, Plcg2*M28L, Ceacam1 KO* (Figure 3C, Supplementary Table S2). Overall, we observed that multiple AD-associated pathways were upregulated in the presence of some LOAD risk variants but downregulated in presence of another set of risk variants. This suggest that distinct risk variants perturb distinct molecular changes associated with LOAD in aging mice.

### Age-dependent pathway effects driving AMP-AD module correlations in *ABCA7*, *MTHFR*, *SORL1*, and *PLCG2* mouse models

In our mouse-human correlation analysis, the effects of multiple LOAD variants (*Abca7*A1527G, Mthfr*677C>T, Sorl1*A528T*, and *Plcg2*M28L*) correlated with human AMP-AD co-expression modules in age-dependent and pathway-specific manner. To further identify the AD-relevant biological processes associated with these selected LOAD risk variants (*Abca7*A1527G, Mthfr*677C>T, Sorl1*A528T*, and *Plcg2*M28L*) we adopted two approaches. First, we performed GSEA (52) on the NanoString Mouse AD Panel genes ranked based on regression coefficients calculated for each factor at 12 months and identified significantly enriched Gene Ontology terms (padj < 0.05). Next, we isolated the homologous genes exhibiting directional coherence between the effects of selected genetic risk variants (*Abca7*A1527G, Mthfr*677C>T, Sorl1*A528T,* and *Plcg2*M28L*) and changes in expression in human AMP-AD co-expression modules at 12 months and performed Gene Ontology (GO) enrichment analysis. These subsets represent the pathways that (1) are altered in each mouse model, and (2) drive the mouse-human module associations. GO terms common to both enrichment tests were then annotated to the modules in which they appear.

The *Abca7*A1527G* variant showed significant negative correlations (p < 0.05) with immune-related modules in Consensus Cluster B, cell cycle and myelination-associated modules in Consensus Cluster D, and cellular stress-response associated modules in Consensus Cluster E (Figure 4A) at four months. However, at 12 months these effects were reversed and the variant exhibited significant positive correlations (p < 0.05) with several immune-related modules in Consensus Cluster B, cell cycle and myelination-associated modules in Consensus Cluster D, and cellular stress-response associated modules in Consensus Cluster E (Figure 4A). Biological processes such as ’de novo’ protein folding, ’de novo’ post-translational protein folding, granulocyte migration, cytokine-mediated signaling pathway, insulin receptor signaling pathway, and neutrophil migration had increased expression in the presence of *Abca7*A1527G* (Figure 4A, Supplementary Table S3). The correlation between the *Abca7*A1527G* variant and the immune-associated human co-expression modules (Consensus Cluster B) (Figure 4A, Supplementary Table S5) was driven by genes enriched for granulocyte migration, cytokine-mediated signaling pathway, and neutrophil migration (including *Pecam1, Cd74, Trem2, Trem1, Csf1*, *Il1rap*, and *Ceacam1*) (Supplementary Table S4). As drivers of the correlations between *Abca7*A1527G* and Consensus Cluster E modules (Figure 4A, Supplementary Table S5), we found genes enriched in ‘de novo’ protein folding and ’de novo’ post-translational protein folding (*e.g., Hspa2*, *Hspa1b*, and *Dnajb4*) (Supplementary Table S4). Insulin receptor signaling was enriched in genes (*Foxo1, Prkcq,* and *Bcar3*) (Supplementary Table S4) driving the correlation between *Abca7*A1527G* and modules in Consensus Cluster D (Figure 4A, Supplementary Table S5).

**FIGURE 4:**
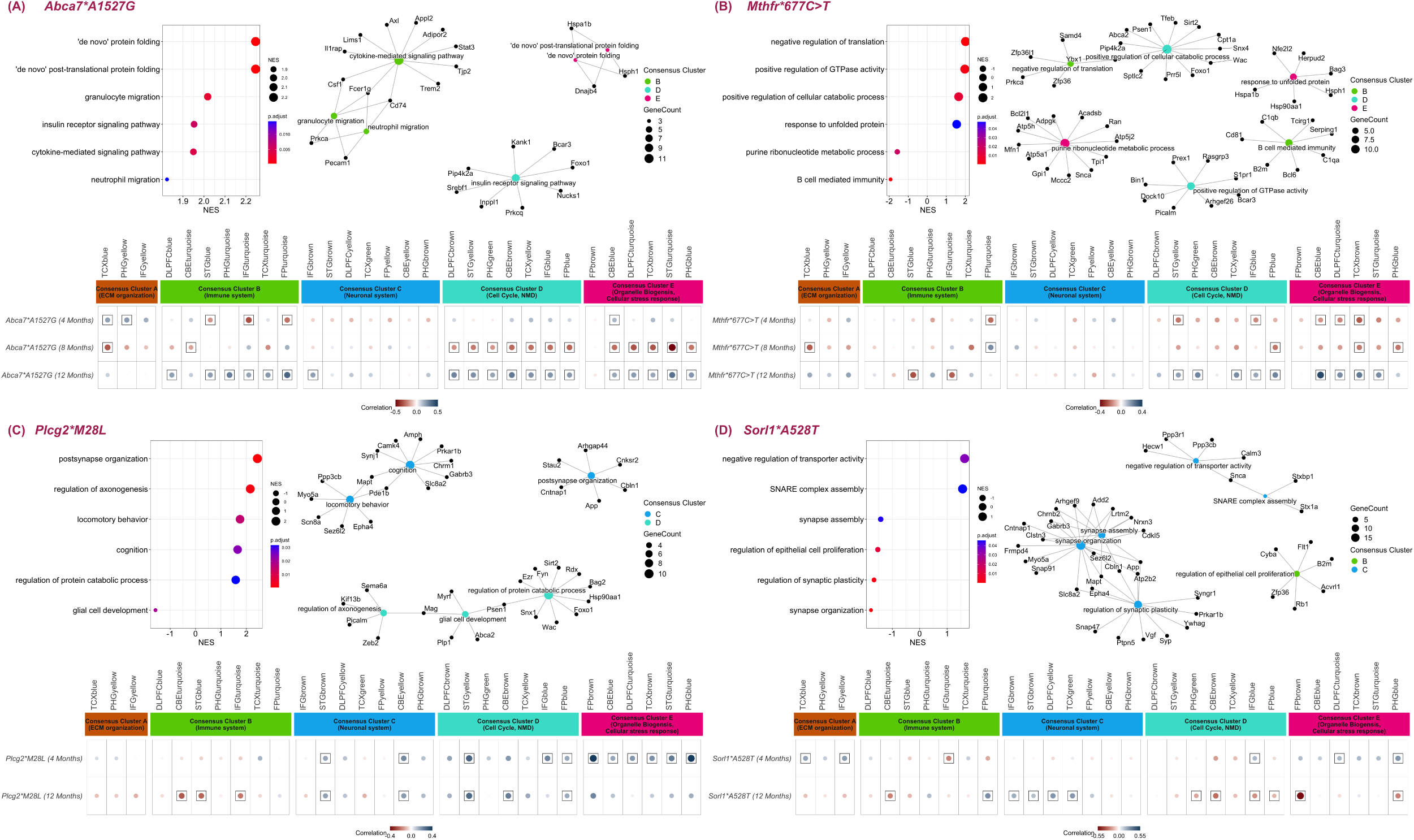
Identification of specific AD-associated processes in LOAD risk variants exhibiting transcriptomic changes similar to human LOAD in age-dependent manner. For four new mouse strains the following is displayed: the six top enriched GO terms identified by GSEA and GO enrichment analysis of genes with common directional changes with human AD modules (top left); gene module networks with common directional changes with the human AMP-AD modules, where node colors correspond to human AMP-AD Consensus Clusters A (orange), B (green), C (blue), D (turquoise) or E (pink) (top right); and the effects of each variant at multiple ages correlated across LOAD effects in 30 AMP-AD modules, following the legend of Figure 3. **(A)** results for the *Abca7*A1527G* model, **(B)** results for the *Mthfr*677C>T* model, **(C)** results for the *Plcg2*M28L* model, and **(D)** results for the *Sorl1*A528T* model. All results are relative to the LOAD1 genetic background for all strains.

A similar reversal of effects with age was observed for *MTHFR*. The *Mthfr*677C>T* variants exhibited significant negative correlations (p < 0.05,) with several cell cycle and myelination-associated modules in Consensus Cluster D and cellular stress-response associated modules in Consensus Cluster E (Figure 4B) at four months. At 12 months, these correlations were positive (Figure 4B). GSEA of the *Mthfr*677C>T* variant identified significant enrichments of response to unfolded protein, positive regulation of cellular catabolic process, negative regulation translation, positive regulation of GTPase activity, B cell mediated immunity, and purine ribonucleotide metabolic process (Figure 4B, Supplementary Table S3). B cell mediated immunity and negative regulation translation biological processes were also enriched in genes (including *C1qa, C1qb, Cd81*, and *Zfp36*) (Supplementary Table S4) with directional coherence for *Mthfr*677C>T* and LOAD effects in Consensus Cluster B (Figure 4B, Supplementary Table S5). Correlations between the *Mthfr*677C>T* variant and Consensus Cluster D changes (Figure 4B, Supplementary Table S5) were driven by genes enriched for positive regulation of cellular catabolic process and positive regulation of GTPase activity (including *Bin1, Picalm, Dock10*, and *Psen1*) (Supplementary Table S4). Biological processes such as response to unfolded protein and purine ribonucleotide metabolic process were enriched in genes (*e.g*., *Hspa1b, Hsph1, Hsp90aa1*, *Snca*, and *Atpp5h*) (Supplementary Table S4) underlying the correlations between *Mthfr*677C>T* and Consensus Cluster E effects (Figure 4B, Supplementary Table S5).

The *Plcg2*M28L* variant caused significant positive correlations (p < 0.05) with neuronal-related modules in Consensus Cluster C and cell-cycle associated modules in Consensus Cluster D at both four and 12 months (Figure 4C). Enriched biological processes included postsynapse organization, regulation of axonogenesis, cognition, locomotory behavior, glial cell development, and regulation of protein catabolic process (Figure 4C, Supplementary Table S3). Biological processes such as postsynapse organization, cognition, and locomotory behavior were enriched in genes (*Mapt, Gabrb3, App, Ppp3cb*, and *Slc8a2*) (Supplementary Table S4) with directional coherence for *Plcg2*M28L* human AD changes in Consensus Cluster C (Figure 4C, Supplementary Table S5). Biological processes such as regulation of axonogenesis, glial cell development and regulation of protein catabolic process were enriched in genes (*Snx1, Picalm, Psen1, Mag, Foxo1*, and *Kif13b*) (Supplementary Table S4) drove the correlations between *Plcg2*M28L* and Consensus Cluster D effects (Figure 4C, Supplementary Table S5).

Aged *Sorl1*A528T* mice (12 months) showed positive correlations (p < 0.05) with neuronal-associated modules in Consensus Cluster C that were not apparent at four months of age (Figure 4D). Enriched processes included the downregulation of synapse organization, synapse assembly, regulation of synaptic plasticity and regulation of epithelial cell proliferation, and the increased expression of negative regulation of transporter activity and SNARE complex assembly genes. These processes drove the correlation between the *SORL1* variant and LOAD effects in Consensus Cluster C modules (Figure 4D, Supplementary Table S5), where GSEA for genes with directional coherence generated synapse organization, synapse assembly, regulation of synaptic plasticity, upregulation of negative regulation of transporter activity, and SNARE complex assembly (including the genes *Mapt, App, Gabrb3, Calm3, Snca, Cdkl5, Vgf*, and *Ywhag*) (Supplementary Table S4).

Overall, we found that late-onset genetic factors in mice generally led to both more abundant changes with age and increasingly disease-relevant pathway changes with age.

### Alignment of mouse models with AD Subtypes

Postmortem transcriptomics from AMP-AD and similar studies have enabled the partitioning of AD cases into potential disease subtypes. These studies have often stratified AD subjects into inflammatory and non-inflammatory subtypes (50, 55, 56). To determine if our mouse models preferentially resembled putative AD subtypes, we correlated the effect of each variant with inflammatory and non-inflammatory subtypes associated with LOAD (50) in the ROSMAP, MSBB, and Mayo cohorts (47-49).

We found that at four months of age, variants did stratify by human subtypes. The effects of *Abca7*A1527G, Sorl1*A528T,* and *Plcg2*M28L* were positively correlated (p < 0.05) with the inflammatory subtypes across all three cohorts, while *Mtmr4*V297G* was positively correlated (p < 0.05) with ROSMAP and MSBB inflammatory subtypes (Figure 5). In contrast, *Shc2*V433M* and *Clasp2*L163P* exhibited significant positive correlations (p < 0.05) with non-inflammatory subtypes across all three cohorts (Figure 5).

**FIGURE 5:**
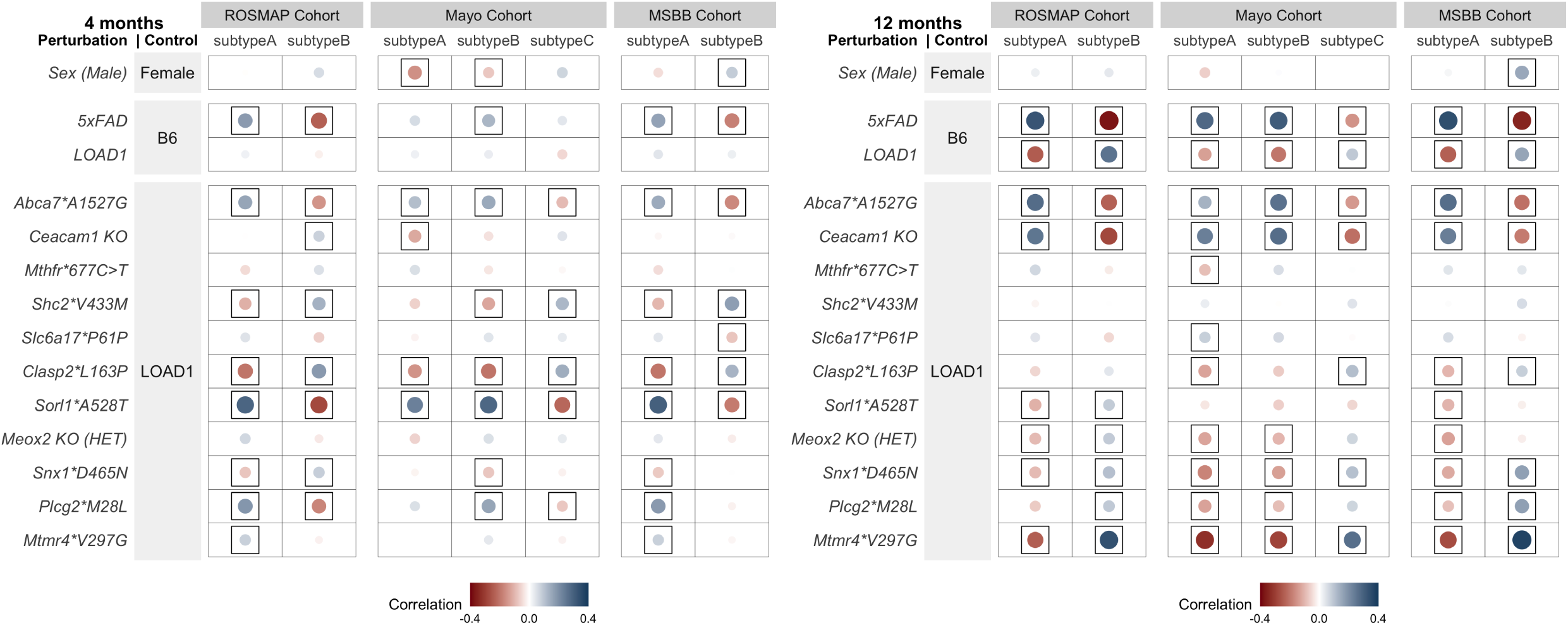
Correlation between the effect of each mouse perturbation and molecular subtypes of LOAD. Two molecular LOAD subtypes inferred in the ROSMAP cohort, three subtypes in the Mayo cohort, and two subtypes in the Mount Sinai Brain Bank (MSBB) cohort (50). Framed circles correspond to significant (p < 0.05) positive (blue) and negative (red) Pearson’s correlation coefficients across all genes on the NanoString panel, with color intensity and circle size proportional to the correlation. **(B)** At four months, the *Abca7*A1527G* and *Sorl1*A528T* variants represent inflammatory subtypes of LOAD (Subtypes A) in each of the cohorts, while *Shc2*V433M* and *Clasp2*L163P* variants mimic the non-inflammatory subtypes of LOAD (Subtypes B). **(C)** At 12 months, the *Abca7*A1527G* and *Ceacam1 KO* variants recapitulate inflammatory subtypes of LOAD (Subtypes A), while the *Snx1*D465N, Mtmr4*V297G*, and *LOAD1* variants model non-inflammatory subtypes of LOAD (Subtypes B).

At 12 months, the correlations between *Abca7*A1527G* effects and the inflammatory subtypes across all three cohorts increased (p < 0.05) and the *Ceacam1 KO* variant had become positively correlated (p < 0.05) with the inflammatory subtypes across all three cohorts (Figure 5). On other hand LOAD1, *Meox2 KO (HET),* and *Snx1*D465N* were positive correlated (p < 0.05) with non-inflammatory subtypes across all three cohorts (Figure 5). Three strains, *Sorl1*A528T, Plcg2*M28L,* and *Mtmr4*V297G*, which were positively correlated (p < 0.05) with inflammatory subtypes at four months, transitioned to correlation (p < 0.05) with non-inflammatory subtypes at 12 months (Figure 5). These results are in concordance with our findings that *Abca7*A1527G* was significantly correlated with immune related human modules and were enriched for immune associated biological processes (Figure 4A), while *Sorl1*A528T* and *Plcg2*M28L* variants were significantly correlated with neuronal related human modules and enriched for neuronal associated biological processes (Figure 4C-D). Overall, these findings suggest that different mouse strains may provide better models for distinct AD subtypes, and that risk for these subtypes may be influenced by distinct AD genetic factors.

## Discussion

In this study, we have performed gene expression screening of new knock-in mouse models harboring candidate genetic variants for late-onset Alzheimer’s disease. Our ultimate goal is to provide the research community and therapeutic development programs with improved preclinical models of LOAD, suitable for preclinical testing of therapeutics that target molecular processes contributing to LOAD origins and progression. By basing these models on human genetics, we also provide a preliminary functional characterization of possible disease-relevant effects from the candidate genetic variants.

Notable results include the finding that many AD-related pathways, modules, and processes are affected by the introduction of late-onset variants. However, the changes were not consistent across strains, suggesting that different genetic loci contribute to distinct AD-related dysfunction (Figures 2 and 4). For example, we determined that the *SORL1* risk factor impinges primarily on AD-relevant synaptic gene expression, while the *ABCA7* variant broadly affected non-neuronal gene expression including immune, protein folding, and metabolic pathways. Meanwhile the *PLCG2* variant primarily affected genes that were annotated to behavior, synapses, and glial cells and similarly changed in human LOAD. We note that a transgenic model harboring familial AD mutations in *App* and *Psen1* exhibited different gene expression changes focused on an acute inflammatory response. Finally, the limited effects of variants like *Clasp2*L163P* suggest that the specific variant is not disease-associated, its AD-related effects are not visible in the transcriptome, and/or it does not trigger changes until later age. This diversity of effects across mouse strains provides specific models to study different aspects of AD biology and paves the way for precision preclinical testing of candidate therapeutics that target these pathways.

Preliminary analysis further suggested that the different loci contribute in an age-dependent manner (Figures 2 and 4) and model putative disease subtypes (Figure 5). However, validation of such partitioning of genetic risk is difficult in human studies due to postmortem tissue sampling and limited cohort size for multi-omic data (50). We also found that the gene expression effects of LOAD variant knock-ins generally increased in terms of magnitude and disease relevance as mice aged from four to 12 months (Figures 2 and 4). This finding supports the notion that LOAD genetic factors become more relevant in an aging brain as they contribute to late-life disease risk.

We note that genetic variants from frequently associated loci tended to produce the most consistent AD-relevant phenotypes (e.g. SORL1, ABCA7, PLCG2) although many of the more exploratory variants also generated AD-like expression signatures across multiple modules in aging mice (e.g. CEACAM1, MTMR4) (Figure 2). Recent advances in variant inference and functional prediction, including many noncoding variants and major GWAS loci, will enable the next round of models to address additional GWAS loci without candidate coding variants, such as the *EPHA1* locus (25). Furthermore, many AD-associated loci suffered from insufficient homology in mice (e.g. *MS4A4/MS4A6E*, *INPP5D, CR1*), which will be addressed by ongoing efforts to humanize these relevant regions of the mouse genome (Benzow K, *et al.,* this issue).

This study had several caveats that need to be noted. Most importantly, aging is the strongest risk factor for late-onset AD (57) and it needs to be recognized that mice at 12 months of age are roughly equivalent to humans at 38-47 years of age. Therefore, our transcriptomic comparison to post-mortem AMP-AD clinical samples, while practical, is unrealistic and we are now testing those models that best approximated human transcriptional changes at 12 months to at least 24 months of age (31, 58) (Oblak A, *et al.,* this issue). Likewise, recent studies (as well as our pilot data) have shown that proteomics is a more reliable means to correlate models to disease than transcriptomics (59, 60) (Oblak A, *et al.,* this issue), so we will be using proteomic analysis on prioritized models.

The Trem2*R47H allele in the LOAD1 base model used here has been shown to cause an ∼2-fold decrease in *Trem2* expression (61). However, our analysis technique factors out allele effects individually so that we are confident of our results. We have since created a new model (JAX #33781) that we have shown has normal Trem2 transcript levels and that will replace the allele used here in future projects.

In this study, we have focused on introducing coding variants on a LOAD1 background (20), aged the mice to middle age (12 months), and characterized the animals using a gene expression panel developed for rapid comparison to recent human study results (21). In future work we will extend our approach to model candidate noncoding variants at LOAD genetic loci without strong candidate coding SNPs, humanizing loci and regulatory regions when necessary (Benzow K, *et al.,* this issue). We will breed the most promising variants presented here – *Abca7*A1527G, Sorl1*A528T, Mthfr*677C>T,* and *Plcg2*M28L* – to a genetic background with humanized Aβ peptide (the LOAD2 strain) and age cohorts beyond 18 months to assess additional disease-related progression with advanced age. These select strains will be assessed in depth with multiple genome-scale omics measures (RNA-seq, tandem mass tag proteomics, metabolomics), plasma biomarkers, *in vivo* imaging, neuropathology and behavioral metrics. Each assay will be optimized for translational value. We will also introduce modifiable risk factors through unhealthy diets and exposure to common environmental toxicants. At the same time, all models are distributed without use restrictions to enable all researchers to obtain, study, and modify these models as desired.

## Acknowledgements

We gratefully acknowledge the contribution of Genome Technology core and Candice Baker and Kim Martens in the Genetic Engineering Technologies Service at The Jackson Laboratory for expert assistance with the work described in this publication. The results published here are in whole or in part based on data obtained from the AD Knowledge Portal (https://adknowledgeportal.org). Study data were provided by the Rush Alzheimer’s Disease Center, Rush University Medical Center, Chicago. Data collection was supported through funding by NIA grants P30AG10161 (ROS), R01AG15819 (ROSMAP; genomics and RNAseq), R01AG17917 (MAP), R01AG36836 (RNAseq), the Illinois Department of Public Health (ROSMAP), and the Translational Genomics Research Institute (genomic). Additional phenotypic data can be requested at www.radc.rush.edu. Mount Sinai Brain Bank data were generated from postmortem brain tissue collected through the Mount Sinai VA Medical Center Brain Bank and were provided by Dr. Eric Schadt from Mount Sinai School of Medicine. The Mayo RNAseq study data was led by Nilüfer Ertekin-Taner, Mayo Clinic, Jacksonville, FL as part of the multi-PI U01 AG046139 (MPIs Golde, Ertekin-Taner, Younkin, Price). Samples were provided from the following sources: The Mayo Clinic Brain Bank. Data collection was supported through funding by NIA grants P50 AG016574, R01 AG032990, U01 AG046139, R01 AG018023, U01 AG006576, U01 AG006786, R01 AG025711, R01 AG017216, R01 AG003949, NINDS grant R01 NS080820, CurePSP Foundation, and support from Mayo Foundation. Study data includes samples collected through the Sun Health Research Institute Brain and Body Donation Program of Sun City, Arizona. The Brain and Body Donation Program was supported by the National Institute of Neurological Disorders and Stroke (U24 NS072026 National Brain and Tissue Resource for Parkinson’s Disease and Related Disorders), the National Institute on Aging (P30 AG19610 Arizona Alzheimer’s Disease Core Center), the Arizona Department of Health Services (contract 211002, Arizona Alzheimer’s Research Center), the Arizona Biomedical Research Commission (contracts 4001, 0011, 05-901 and 1001 to the Arizona Parkinson’s Disease Consortium), and the Michael J. Fox Foundation for Parkinson’s Research.

## Consent Statement

No consent was required as all human subjects data were reused under controlled access with an active AD Knowledge Portal Data Use Certificate (v7.3). All data were anonymized by original sources with no possibility of deanonymization.

## Ethics Statement

The animal study was reviewed and approved by the The Jackson Laboratory Animal Use Committee.

## Availability of data and materials

The MODEL-AD data sets are available via the AD Knowledge Portal (https://adknowledgeportal.org). The AD Knowledge Portal is a platform for accessing data, analyses, and tools generated by the Accelerating Medicines Partnership (AMP-AD) Target Discovery Program and other National Institute on Aging (NIA)-supported programs to enable open-science practices and accelerate translational learning. The data, analyses and tools are shared early in the research cycle without a publication embargo on secondary use. Data is available for general research use according to the following requirements for data access and data attribution (https://adknowledgeportal.org/DataAccess/Instructions).

All mouse models are available from the Jackson Laboratory mouse repository.

## Figure and table captions

**SUPPLEMENTAL FIGURE 1:** Validation of novel mouse models. RNA-seq was performed on brain tissue at 4 months of age for each model. **(A)** Sequence analysis identified appropriate engineered variants; in some cases, silent mutations were introduced for CRISPR or genotyping purposes. **(B)** Transcript counts were used to demonstrate normal expression levels for SNP models, and lack of expression in the Ceacam1 knockout model.

**SUPPLEMENTAL TABLE 1:** Reagents used to engineer LOAD mutations using CRISPR. The Meox2 allele was previously created (62) and obtained from the JAX repository (JAX # 3755).

**SUPPLEMENTAL TABLE 2:** Gene set enrichment analyses results of Reactome pathways for the effects of LOAD risk variants in mice at 12 months.

**SUPPLEMENTAL TABLE 3:** Gene set enrichment analyses results of GO terms for the effects of Abca7*A1527G, *Mthfr*677C>T, Plcg2*M28L, and Sorl1*A528T* variants in mice.

**SUPPLEMENTAL TABLE 4:** Genes with common directional changes for the effects of Abca7*A1527G, *Mthfr*677C>T, Plcg2*M28L, and Sorl1*A528T* variants in mice at 12 months and human AD cases.

**SUPPLEMENTAL TABLE 5:** Enriched GO terms in genes with common directional changes for the effects of Abca7*A1527G, *Mthfr*677C>T, Plcg2*M28L, and Sorl1*A528T* variants in mice at 12 months and human AD cases.

